# Bioinformatics Core Competencies for Undergraduate Life Sciences Education

**DOI:** 10.1101/170993

**Authors:** Melissa A. Wilson Sayres, Charles Hauser, Michael Sierk, Srebrenka Robic, Anne G. Rosenwald, Todd M. Smith, Eric W. Triplett, Jason J. Williams, Elizabeth Dinsdale, William Morgan, James M. Burnette, Samuel S. Donovan, Jennifer C. Drew, Sarah C. R. Elgin, Edison R. Fowlks, Sebastian Galindo-Gonzalez, Anya L. Goodman, Neal F. Grandgenett, Carlos C. Goller, John Jungck, Jeffrey D. Newman, William Pearson, Elizabeth Ryder, Rafael Tosado-Acevedo, William Tapprich, Tammy C. Tobin, Arlín Toro-Martínez, Lonnie R. Welch, Robin Wright, David Ebenbach, Kimberly C. Olney, Mindy McWilliams, Mark A. Pauley

## Abstract

Bioinformatics is becoming increasingly central to research in the life sciences. However, despite its importance, bioinformatics skills and knowledge are not well integrated in undergraduate biology education. This curricular gap prevents biology students from harnessing the full potential of their education, limiting their career opportunities and slowing genomic research innovation. To advance the integration of bioinformatics into life sciences education, a framework of core bioinformatics competencies is needed. To that end, we here report the results of a survey of life sciences faculty in the United States about teaching bioinformatics to undergraduate life scientists. Responses were received from 1,260 faculty representing institutions in all fifty states with a combined capacity to educate hundreds of thousands of students every year. Results indicate strong, widespread agreement that bioinformatics knowledge and skills are critical for undergraduate life scientists, as well as considerable agreement about which skills are necessary. Perceptions of the importance of some skills varied with the respondent’s degree of training, time since degree earned, and/or the Carnegie classification of the respondent’s institution. To assess which skills are currently being taught, we analyzed syllabi of courses with bioinformatics content submitted by survey respondents. Finally, we used the survey results, the analysis of syllabi, and our collective research and teaching expertise to develop a set of bioinformatics core competencies for undergraduate life sciences students. These core competencies are intended to serve as a guide for institutions as they work to integrate bioinformatics into their life sciences curricula.

**Significance Statement:** Bioinformatics, an interdisciplinary field that uses techniques from computer science and mathematics to store, manage, and analyze biological data, is becoming increasingly central to modern biology research. Given the widespread use of bioinformatics and its impacts on societal problem-solving (e.g., in healthcare, agriculture, and natural resources management), there is a growing need for the integration of bioinformatics competencies into undergraduate life sciences education. Here, we present a set of bioinformatics core competencies for undergraduate life scientists developed using the results of a large national survey and the expertise of our working group of bioinformaticians and educators. We also present results from the survey on the importance of bioinformatics skills and the current state of integration of bioinformatics into biology education.

## Introduction

Over the past two decades, the rapid development of high-throughput technologies, data storage capacity, and sophisticated algorithms has produced substantial changes in research practices in the life sciences and medicine. In order for researchers and practitioners in these areas to take advantage of the changes in data availability, they need to have computational and quantitative skills—such as those encompassed by bioinformatics—beyond what was required of them in the past. For more than a decade, authoritative calls from a variety of professional organizations to update undergraduate life sciences curricula have stressed the importance of increasing quantitative and computational education to prepare life sciences students for 21st-century careers (1–14). Examples include *BIO2010: Transforming Undergraduate Education for Future Research Biologists*, a report by the National Academy of Sciences (1); *Math & Bio 2010: Linking Undergraduate Disciplines*, a joint project of the American Association for the Advancement of Sciences, the American Society for Mathematical Biology, and the Mathematical Association of America (11); *Scientific Foundations for Future Physicians*, a publication from the Association of American Medical Colleges and the Howard Hughes Medical Institute (12); and *Vision and Change in Undergraduate Biology Education: A Call to Action* (13) and *Chronicling Change, Inspiring the Future* (14). These reports are consistent in their recommendations that life sciences majors should receive better training in chemistry, physics, mathematics, and computation because such knowledge and skills are necessary to address questions at all levels of biology. In addition, several publications have called for life scientists to develop more sophisticated data analysis and programming skills so that they can benefit from the bioinformatics revolution (7, 15–20). These include a report from the RECOMB Bioinformatics Education Conference (7) and a recent essay in *Nature* (17). Along these lines, it is important to note that in its most recent employment predictions, the United States Bureau of Labor Statistics projected that between now and 2024, 75% of new science, technology, engineering, and mathematics (STEM) jobs will involve computation (21,22).

On a national scale, a number of innovative programs have been created to address the need for increased quantitative and computational education, including the Science Education Alliance – Phage Hunters Advancing Genomics and Evolutionary Science (SEA-PHAGES) program (23), Genomics Education Partnership (GEP) (3,24), Genome Consortium for Active Teaching (GCAT) (4), GCAT NextGen Sequencing Group (GCAT-SEEK) (25), and Genome Solver (10). On a local scale, we are aware that the number of courses and meetings focused on undergraduate bioinformatics is increasing and that many existing courses have been modified to include bioinformatics. However, despite these efforts, the adoption of bioinformatics is often limited either to a small number of institutions or to particular courses within a given biology curriculum. As an example, SEA-PHAGES engages first-year undergraduates in a genuine research experience and includes substantial bioinformatics analysis as part of its year-long curriculum. As reported in 2014, the SEA-PHAGES program had been used at seventy-three institutions by 4,800 students over five years (23). As impressive as the reach of SEA-PHAGES has been, this program impacts only a small fraction of the thousands of US institutions that offer a biology degree and the approximately 110,000 students who now graduate annually with biological and biomedical sciences degrees (not to mention the 220,000 more who graduate from health professions and related programs) (26). In addition, because the SEA-PHAGES program is designed for first-year students, it does not necessarily impact the rest of the curriculum at a particular institution. The reaches of the other mentioned programs are similar to, or not as extensive as, that of SEA-PHAGES. Despite these large-scale efforts and those of independent faculty to build resources for bioinformatics education, bioinformatics is not yet a standard component of life sciences education (15).

Several groups have made independent efforts to develop or describe bioinformatics curricula, but they have primarily focused on training undergraduate and graduate bioinformaticians, not undergraduate life scientists. For example, the Curriculum Task Force of the Education Committee of the International Society for Computational Biology (ISCB) surveyed directors of bioinformatics core facilities in Europe, Israel, the United States, and Canada about the skills needed for success in the field of bioinformatics and what skills were lacking in the bioinformaticians they recently hired. Based on the results, the Task Force developed a set of core competencies for bioinformatics professionals (9,27). They also described three professional roles that require bioinformatics training and the different but overlapping competencies required for individuals in those roles. As another example, Koch and Fuellen described the bioinformatics curricula at various German universities at both the graduate and undergraduate level (28). However, a few authors have considered the specific bioinformatics needs of general life scientists. For example, Maloney et al. discussed the importance of incorporating bioinformatics into undergraduate biology curricula and presented examples of where this has been done successfully (29). The East Asia Bioinformation Network developed a bioinformatics skill set for biology students in the developing countries that are part of the Association for Southeast Asian Nations (https://eabn.apbionet.org/3eabn08/docs.shtml). Finally, Tan et al. proposed a minimum bioinformatics skill set for life sciences curricula (graduate and undergraduate) in developed nations, formulated from discussions at a conference focused on education in the Asia-Pacific region (8). Thanks to developments such as these, the state of integration of bioinformatics into life sciences education is maturing.

The Network for Integrating Bioinformatics into Life Sciences Education (NIBLSE, pronounced “nibbles”) is using an evidence-based approach to expand and promote the integration of bioinformatics skills and knowledge into undergraduate life sciences education (30). NIBLSE is a National Science Foundation (NSF) Undergraduate Biology Education Research Coordination Network (RCN-UBE) that was formed in 2014 to build on and expand the curricular developments mentioned above. Since further integration of bioinformatics into undergraduate life sciences education requires the participation of non-expert faculty, one of the goals of NIBLSE is to provide resources (e.g., curricular materials and assessments) and support to faculty interested in expanding the integration of bioinformatics within their departments and programs. Note that NIBLSE is focused on the integration of bioinformatics into undergraduate life sciences curricula, which is related to, but distinct from, the curricular requirements for a “pure” bioinformatics degree. The goals of NIBLSE and the ISCB Task Force mentioned above are thus different.

In order for NIBLSE to provide curricular resources and assessment tools that align with the needs of undergraduate life sciences students, a set of core bioinformatics competencies for the training of these students is needed as a framework. To that end, NIBLSE recently conducted a survey (hereafter the “NIBLSE survey” or just the “survey”) that targeted US biologists. The survey was designed to assess the importance of bioinformatics and bioinformatics skills for undergraduate life scientists, and it included questions about teaching bioinformatics to this group of students and the challenges involved in doing so. In addition, to get a sense of which bioinformatics skills faculty are covering now, respondents were asked to submit syllabi from their courses that incorporate bioinformatics. The analyses of the survey results and syllabi were the subjects of a national NIBLSE conference in August 2016. Outcomes of the analyses and conference are described below and include a proposed set of core bioinformatics competencies for undergraduate life sciences majors. A separate paper now in preparation addresses the barriers to integrating bioinformatics into life sciences education and NIBLSE’s recommendations for overcoming them.

The NIBLSE Bioinformatics Core Competencies for Undergraduate Life Scientists proposed here are intended to serve as a guide for institutions and departments as they integrate bioinformatics into their own life sciences curricula. To facilitate this necessary and important change, NIBLSE is actively collecting curricular resources for faculty and will soon be developing assessment tools and faculty development resources that are aligned with the core competencies.

## Materials and Methods

### Survey development and dissemination

The NIBLSE Core Competencies Working Group (CCWG), composed of biologists and bioinformaticians from a range of educational institutions and industry, developed the survey using an iterative process over the course of several months. Feedback from other NIBLSE members and two evaluation specialists, one with expertise in STEM education, was used to improve the questions and layout of the survey. The survey was implemented and distributed using Qualtrics (https://www.qualtrics.com) with assistance from the Center for New Designs in Learning and Scholarship at Georgetown University. Approval for the study was obtained from the University of Nebraska at Omaha Institutional Review Board (IRB # 161-16-EX) before it was distributed.

The survey was branched, with some questions or sections presented or skipped based on responses to filtering questions. For example, the survey branched depending on whether the respondent taught at a four-year institution, a two-year institution, or provided not-for-credit training (e.g., for a company or organization). The branched structure allowed us to formulate targeted questions. The survey in its entirety is provided as a supplementary document.

One section of the survey asked respondents to rate fifteen bioinformatics skills (S1 to S15, hereafter the “survey skills”) using a five-level Likert scale. The survey skills are listed below; the text in parentheses is the abbreviation of that skill.

- S1 (*Role*)—Understand the role of computation and data mining in hypothesis-driven processes within the life sciences
- S2 (*Concepts*)—Understand computational concepts used in bioinformatics, e.g., meaning of algorithm, bioinformatics file formats
- S3 (*Statistics*)—Know statistical concepts used in bioinformatics, e.g., e-value, z-scores, t test, type-1 error, type-2 error, employ R
- S4 (*Access genomic*)—Know how to access genomic data, e.g., in NCBI nucleotide databases
- S5 (*Tools genomic*)—Be able to use bioinformatics tools to analyze genomic data, e.g., BLASTN, genome browser
- S6 (*Access expression*)—Know how to access gene expression data, e.g., in UniGene, GEO, SRA
- S7 (*Tools expression*)—Be able to use bioinformatics tools to analyze gene expression data, e.g., GeneSifter, David, ORF Finder
- S8 (*Access proteomic*)—Know how to access proteomic data, e.g., in NCBI protein databases
- S9 (*Tools proteomic*)—Be able to use bioinformatics tools to examine protein structure and function, e.g., BLASTP, Cn3D, PyMol
- S10 (*Access metabolomic*)—Know how to access metabolomic and systems biology data, e.g., in the Human Metabolome Database
- S11 (*Pathways*)—Be able to use bioinformatics tools to examine the flow of molecules within pathways/networks, e.g., Gene Ontology, KEGG
- S12 (*Metagenomics*)—Be able to use bioinformatics tools to examine metagenomics data, e.g., MEGA, MUSCLE
- S13 (*Scripting*)—Know how to write short computer programs as part of the scientific discovery process, e.g., write a script to analyze sequence data
- S14 (*Software*)—Be able to use software packages to manipulate and analyze bioinformatics data, e.g., Geneious, Vector NTI Express, spreadsheets
- S15 (*Computational environment*)—Operate in a variety of computational environments to manipulate and analyze bioinformatics data, e.g., Mac OS, Windows, web- or cloud-based, Unix/Linux command line

In addition, a free-response question allowed respondents to specify skills they thought were missing from the list. The skills listed in the survey were based on the *CourseSource* Bioinformatics Learning Framework developed in 2014 by members of NIBLSE and refined with feedback from a number of groups including GEP and GCAT-SEEK (31). (*CourseSource* is an open-access journal of peer-reviewed teaching resources for undergraduate biological sciences.) In turn, the Learning Framework was informed by the core competencies for bioinformaticians developed by the ISCB Curriculum Task Force (9,27). Two members of the CCWG serve on the ISCB Task Force, one of whom serves as co-chair.

A list of more than 11,000 randomly-selected email addresses of biologists at US institutions of higher education was purchased from MDR, an education marketing company (http://schooldata.com). The list included faculty at both four-year and two-year institutions at an approximately 70:30 ratio. Using Qualtrics, unique links to the survey were emailed to the addresses in this list; unique links allowed us to tie a given response to an email address and therefore to the home institution of the respondent. A generic link to the survey was sent to the members of scientific organizations, including the Society for the Advancement of Biology Education Research, GEP, and Biology Scholars alumni (American Society for Microbiology). (Responses to the generic survey link could not be identified by institution.) In addition, the survey was advertised on Twitter and was announced in the CyVerse, Digital World Biology, Bio-Link, American Society for Cell Biology, and National Association of Biology Teachers newsletters as well as on the ScienceBlogs website. Finally, respondents were asked to forward the generic link to colleagues.

### Survey results and analysis

A total of 1,260 responses to the survey were received (*n* = 1,260), 82% of which came from unique links (Fig. S1). Survey results were analyzed by a subset of the NIBLSE CCWG, the interdisciplinary Core Competencies Team (CCT). Differences in Likert responses by self-reported demographics were determined by applying a two-sample Kolmogorov-Smirnov (KS) test to pairwise comparisons of groups. The KS tests were implemented in scripts written in R; these scripts are available from the NIBLSE repository on GitHub (32).

Using information provided to us by MDR and from publicly available databases (33–35), we estimate that there are currently between 75,000 and 100,000 biological sciences faculty in the US. Given our overall sample size (1% to 2%) and the number of responses per question (between 970 and 1,009), we estimate that the mean margin of error for the survey questions described in this paper is ±3% at the 95% confidence interval.

### Syllabi assessment

Ninety syllabi of courses with bioinformatics content were uploaded by survey respondents. Most (sixty-nine) were known or inferred to be from departments of biology, with the balance being from computer science/engineering (nine), biochemistry (five), math (three), and chemistry (two) departments; the home department of two syllabi could not be determined. Each syllabus was assessed independently by two CCT members, who determined which of the fifteen skills (S1 to S15) were covered in the course. When two assessments did not agree (approximately 25% of the time), the discordance was resolved by a third CCT member. Additionally, each member kept track of additional skills (i.e., not one of the S1 to S15 survey skills) covered in each syllabus.

### Development of the core competencies

Using an iterative process, the survey skills were refined into a set of nine core competencies based on results from the survey, comments from survey respondents, assessment of syllabi, and the collective expertise and experience of the authors in both teaching bioinformatics and applying it to research. As mentioned previously, the CCT analyzed the survey results and syllabi with respect to the survey skills. Based on this work, the CCT developed a set of tentative competencies. Included in this set were items that showed up repeatedly in the syllabi and responses to the free-response question but were not in the S1 to S15 survey skills. On the other hand, some of the S1 to S15 survey skills were revised or dropped from the tentative competencies set due to weak support, including S10 (*Access metabolomic*) and S15 (*Computational environment*). Prior to the August 2016 NIBLSE conference, the tentative set of competencies was distributed to the conference participants for their review and consideration.

At the NIBLSE conference, the CCT formally presented the tentative competencies to the conference participants and summarized the evidence on which they were based (survey results, syllabi assessment, etc.). The conference attendees then broke into small groups, each moderated by a member of the CCT. The groups discussed the evidence and revised the competencies based on both their interpretation of the evidence and their own expertise and experience. The groups then reconvened into one large group to discuss the resulting lists and to develop a consensus set of competencies. During this discussion, considerations were made to balance specificity with generalizability. The result of this discussion was a set of nine core competencies agreed to by all of the conference participants. Finally, the conference attendees voted to allow the NIBLSE leadership team (PI and Co-PIs of the NSF RCN-UBE) to decide on the specific final wording of the competencies. The resulting set of competencies, the NIBLSE Bioinformatics Core Competencies for Undergraduate Life Scientists (hereafter the “NIBLSE Core Competencies” or just the “Core Competencies”), is presented in Table 2.

## Results

The NIBLSE CCWG developed and disseminated the survey, with the CCT taking the lead in the analysis of the results and initial development of the NIBLSE Core Competencies (see Materials and Methods).

### NIBLSE survey

The survey assessed Likert-scale responses on the perceived importance of each of fifteen bioinformatics skills (Fig. 1, Fig. 2, and Table 1). We refer to the fifteen skills described in the survey as S1 to S15 and/or by text abbreviation—e.g., S1 (*Role*)—to distinguish them from the nine Core Competencies (C1 to C9) ultimately developed (Table 2). Skills that received the highest mean responses overall included S1 (*Role*), S3 (*Statistics*), S4 (*Access genomic*), and S5 (*Tools genomic*) (Table 1). As detailed below, the perceived importance of some skills varied based on the demographics of the respondents.

**Fig. 1.**
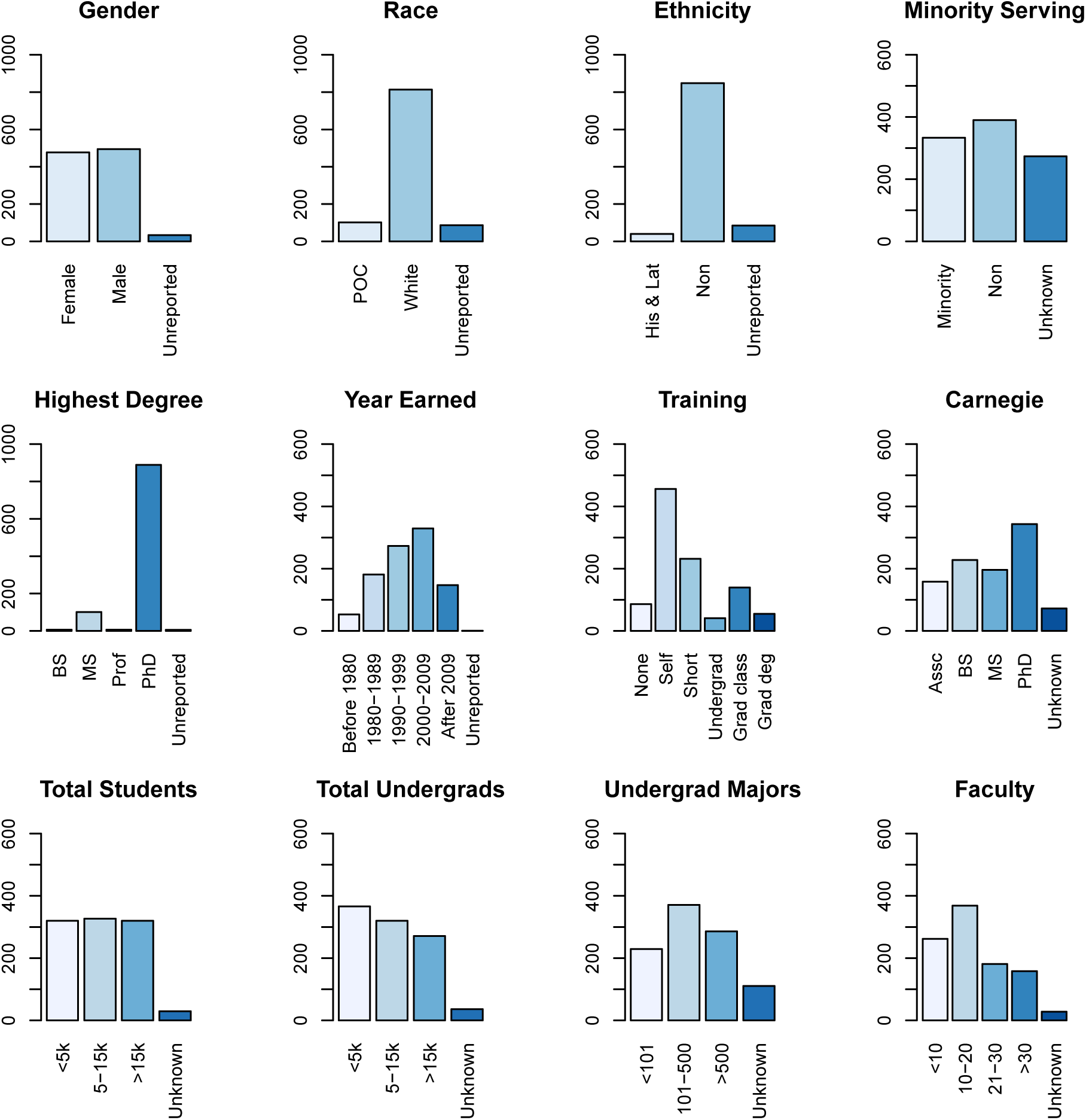
Survey Demographics. The number of responses for the categories of a given demographic variable. The NIBLSE survey asked for information on Gender, Race (People of Color and White), Ethnicity (Hispanic-Latino, non-Hispanic/non-Latino), Minority Serving (whether the respondent’s home institution is classified as minority-serving or not), Highest Degree (highest degree earned: Bachelor’s, Master’s, Professional Degree, PhD), Year Earned (year that the highest degree was earned; responses were grouped in the following bins: Before 1980, 1980 to 1989, 1990 to 1999, 2000 to 2009, and After 2009), Training (level of bioinformatics training: None, Self-taught, Short workshop, Undergraduate/PostBacc training, Graduate class, and Graduate degree), Carnegie (Carnegie classification of the respondent’s home institution: Associate’s, Baccalaureate, Master’s, Doctoral), Total Students (total number of students at the respondent’s home institution), Total Undergraduates (number of undergraduates at the respondent’s home institution), Undergraduate Majors (number of undergraduate majors in the respondent’s home department), and Faculty (number of faculty in the respondent’s home department). Four categories in Training were grouped together into “Undergrad” (undergraduate/ post-baccalaureate training) due to very small sample numbers for each: Undergraduate course (24), Undergraduate certificate (0), Undergraduate degree (5), and Post-baccalaureate certificate (12) for a total of 41 responses.

**Fig. 2.**
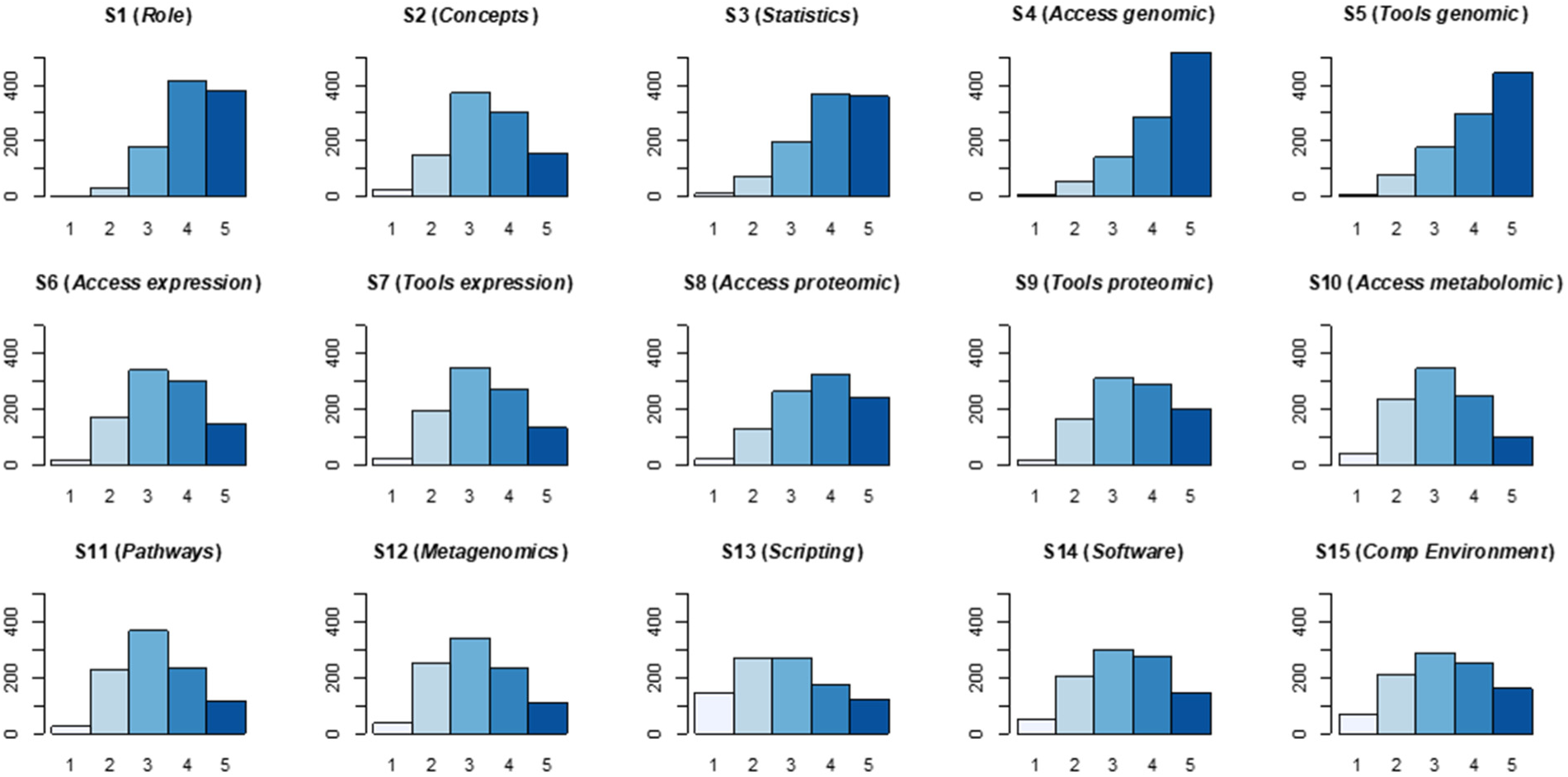
Summary of surveyed skills. The total number of Likert-scale responses from 1 to 5—1 being “Not at all important” to 5 being “Extremely Important”—for each of the fifteen survey skills. As discussed in the narrative, these skills were divided into two broad categories: skills that just required familiarity (“knowing” skills: S1 to S4, S6, S8, S10), and those that required direct engagement (“practicing” skills: S5, S7, S9, S11 to S15).

**Table 1.**
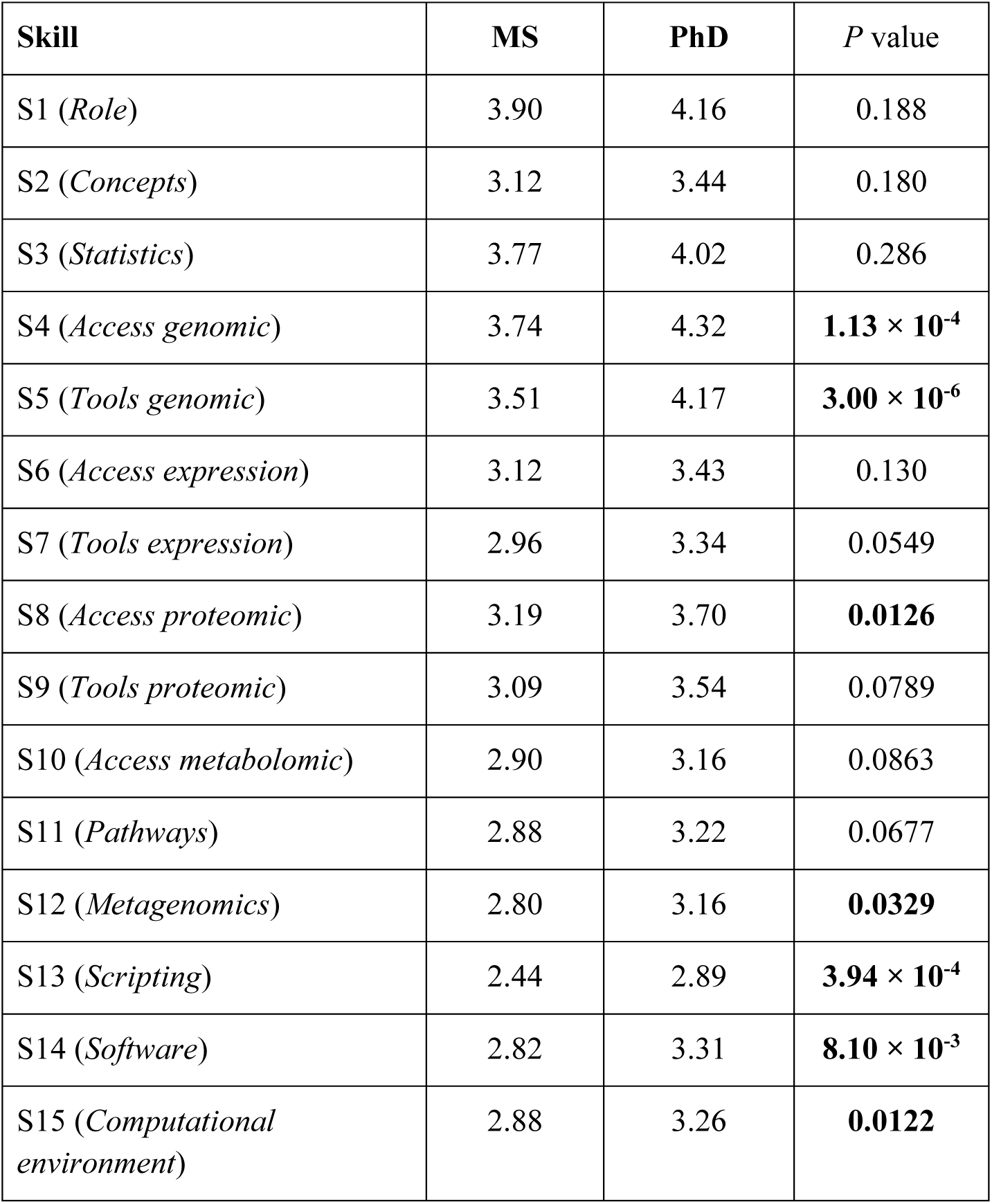
Responses by degree. Means of the Likert-scale responses for each survey skill for respondents with a Master’s degree (MS) and those with a PhD. Two-sided *P* value from a Kolmogorov-Smirnov test of the Likert-scale responses is shown. In all cases, those with a PhD rated desired skills higher than those with an MS; significant differences are in bold.

| Skill | MS | PhD | *P* value |
| --- | --- | --- | --- |
| S1 (*Role*) | 3.90 | 4.16 | 0.188 |
| S2 (*Concepts*) | 3.12 | 3.44 | 0.180 |
| S3 (*Statistics*) | 3.77 | 4.02 | 0.286 |
| S4 (*Access genomic*) | 3.74 | 4.32 | **$1.13 \times 10^{-4}$** |
| S5 (*Tools genomic*) | 3.51 | 4.17 | **$3.00 \times 10^{-6}$** |
| S6 (*Access expression*) | 3.12 | 3.43 | 0.130 |
| S7 (*Tools expression*) | 2.96 | 3.34 | 0.0549 |
| S8 (*Access proteomic*) | 3.19 | 3.70 | **0.0126** |
| S9 (*Tools proteomic*) | 3.09 | 3.54 | 0.0789 |
| S10 (*Access metabolomic*) | 2.90 | 3.16 | 0.0863 |
| S11 (*Pathways*) | 2.88 | 3.22 | 0.0677 |
| S12 (*Metagenomics*) | 2.80 | 3.16 | **0.0329** |
| S13 (*Scripting*) | 2.44 | 2.89 | **$3.94 \times 10^{-4}$** |
| S14 (*Software*) | 2.82 | 3.31 | **$8.10 \times 10^{-3}$** |
| S15 (*Computational environment*) | 2.88 | 3.26 | **0.0122** |

#### Demographics of survey respondents

The group of survey respondents was split fairly evenly between males and females, across institutions by Carnegie classification, by measurement of institution size (total students, total undergraduates, majors, and faculty), and by self-reported minority-serving status versus not minority-serving (Fig. 1). Respondents were from all fifty states with the distribution of responses roughly matching the population distribution of the country (Fig. S1). The respondents were predominantly white and non-Latino/Hispanic; for the vast majority, a PhD was the highest earned degree (Fig. 1). The level of bioinformatics training varied widely among respondents, including no training, self-taught, short courses, undergraduate and graduate classes, and graduate degrees (Fig. 1).

### Analysis of survey results

The Likert responses on the perceived importance of the survey skills are summarized in Fig. 2. Analysis of survey results focused on determining, using the two-sample KS test, whether the Likert responses differed significantly across any demographic categories.

#### No difference by gender or minority-serving institution status

Analyzing the results for the fifteen survey skills, responses did not vary based on gender or whether the respondent was at a minority-serving institution (see Materials and Methods and [32]). Respondents were overwhelmingly white and non-Hispanic/non-Latino (Fig. 1). Individuals who identified as people of color (POC) and those who identified as Hispanic/Latino consistently rated the importance of every skill higher (and often significantly higher; i.e., indicated that it was more important) than those who did not identify as POC or Hispanic/Latino (32). However, given the tremendous disparity in sample sizes (Fig. 1), and known cultural differences in responses to surveys (36,37), we hesitate to assign much weight to these particular differences.

#### Higher ratings of some skills at larger institutions

Respondents gave some skills different scores depending on the size of their institution, whether that is determined by total number of students, total number of undergraduates, or number of faculty in the department. Respondents at an institution with fewer than 5,000 students or 5,000 undergraduates rated the importance of skills S1 (*Role*), S2 (*Concepts*), S13 (*Scripting*), and S15 (*Computational environment*) significantly lower (i.e., indicated that they were less important) than those at institutions with more than 15,000 students or 15,000 undergraduates (32). Similarly, faculty in larger departments gave significantly higher ratings for skills S1 (*Role*) and S2 (*Concepts*) than those in smaller departments (S1: *p* = 6.73 × 10^-3^ for a two-tailed KS test; S2: *p* = 1.49 × 10^-3^).

Among the four Carnegie classifications, there were no significant differences in Likert ratings between respondents at Baccalaureate and Master’s institutions (Table S1). In contrast, respondents at Associate’s institutions routinely rated every skill lower than did those at other institution types, whereas those at Doctoral institutions rated every skill higher than those at institutions with other classifications. In general, the rating of S13 (*Scripting*) increased with Carnegie classification (Fig. 3 and Table S1).

**Fig. 3.**
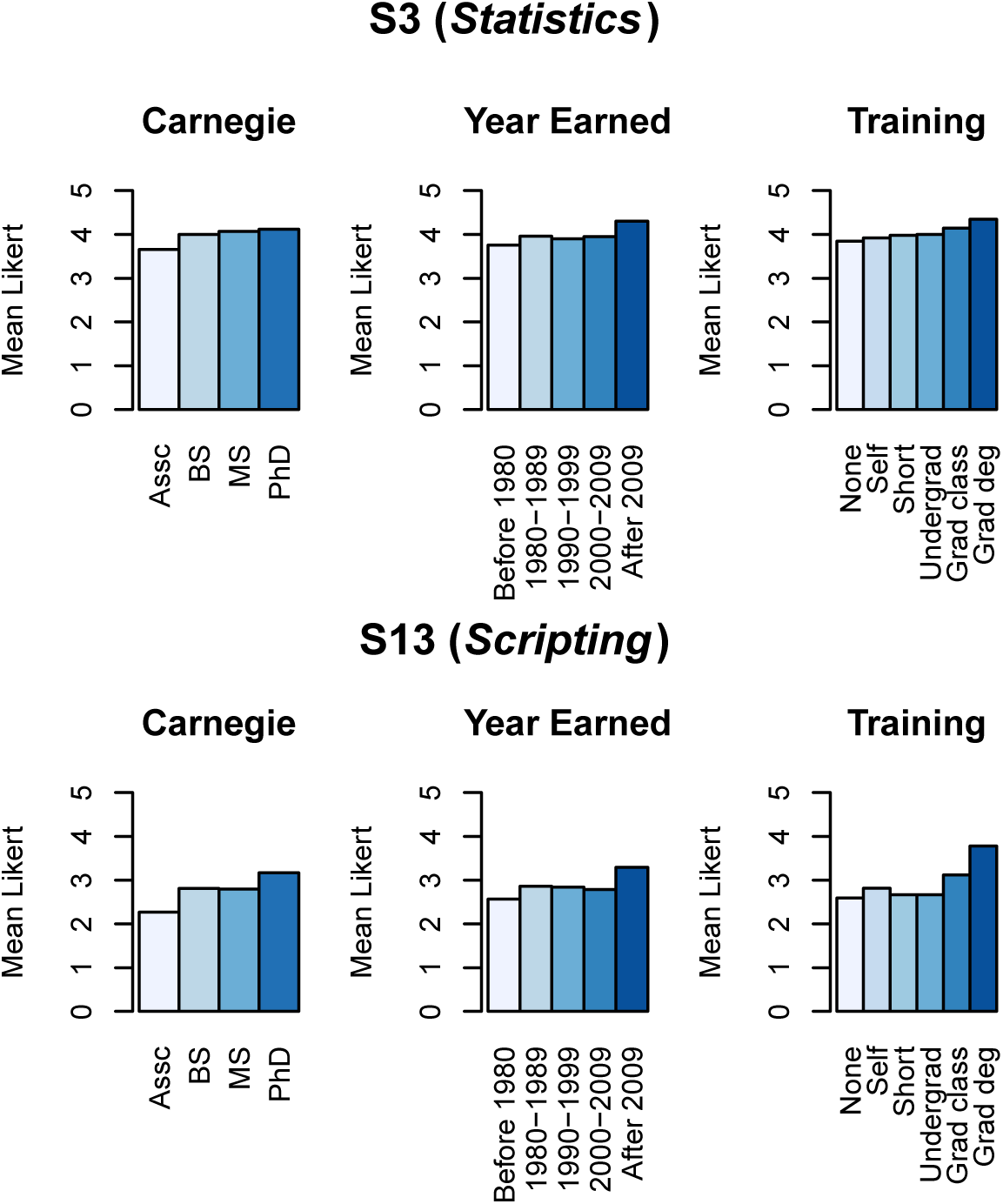
Mean Likert responses for S3 (*Statistics*) and S13 (*Scripting*). Mean Likert responses are shown for S3 (*Statistics*) and S13 (*Scripting*) for three categories: Carnegie (Carnegie classification of the respondent’s home institution: Associate’s, Baccalaureate, Master’s, Doctoral), Year Earned (year that the highest degree was earned; responses were grouped in the following bins: Before 1980, 1980 to 1989, 1990 to 1999, 2000 to 2009, and After 2009), and Training (level of bioinformatics training: None, Self-taught, Short workshop, Undergraduate/PostBacc training, Graduate class, and Graduate degree). Means and *P* values from pairwise KS tests are reported in (32).

#### Higher ratings of categories by year and level of bioinformatics training

The ratings of almost all of the fifteen skills did not depend significantly on the year in which the respondents earned their degree. However, skills S3 (*Statistics*) and S13 (*Scripting*) were rated significantly higher by respondents who earned their degree in 2010 or after than by those who earned their degree prior to 2010 (Fig. 3 and [32]). Furthermore, the majority of respondents had a PhD, and those with a PhD consistently rated every skill higher than those with a master’s degree, often significantly higher (Table 1). Those with no training in bioinformatics routinely rated every skill significantly lower than respondents with training, formal or not (Fig. 3 and [32]). Notably, S13 (*Scripting*) was rated significantly higher by respondents with a graduate degree in bioinformatics than by those with any other type of training (Fig. 3 and [32]).

#### Coverage of skills varies across syllabi

To analyze the ninety syllabi submitted by survey respondents (see Materials and Methods), the fifteen survey skills were divided into two groups—those that required students to be familiar with a concept (i.e., to “know” about it; survey skills S1 to S4, S6, S8, and S10), and those that required direct engagement (i.e., to be able to use the skill in “practice”; skills S5, S7, S9, and S11 to S15). More of the “knowing” skills were rated as being either “extremely important” or “very important” than the “practice” skills and were more frequently covered in the syllabi; conversely, the “practice” skills were less likely to be covered in syllabi (Fig. 4). There were two exceptions to these trends. Few survey respondents thought students should be expected to be familiar with metabolomic and systems biology data (S10, a “knowing” skill), nor was it frequently covered in the submitted syllabi. (As mentioned above, this skill was dropped from the Core Competencies.) On the other hand, a majority of respondents indicated that undergraduate life scientists should “be able to use bioinformatics tools” (S5). In addition, evidence of this was present in approximately 70% of the submitted syllabi (Fig. 4, Fig. S2).

**Fig. 4.**
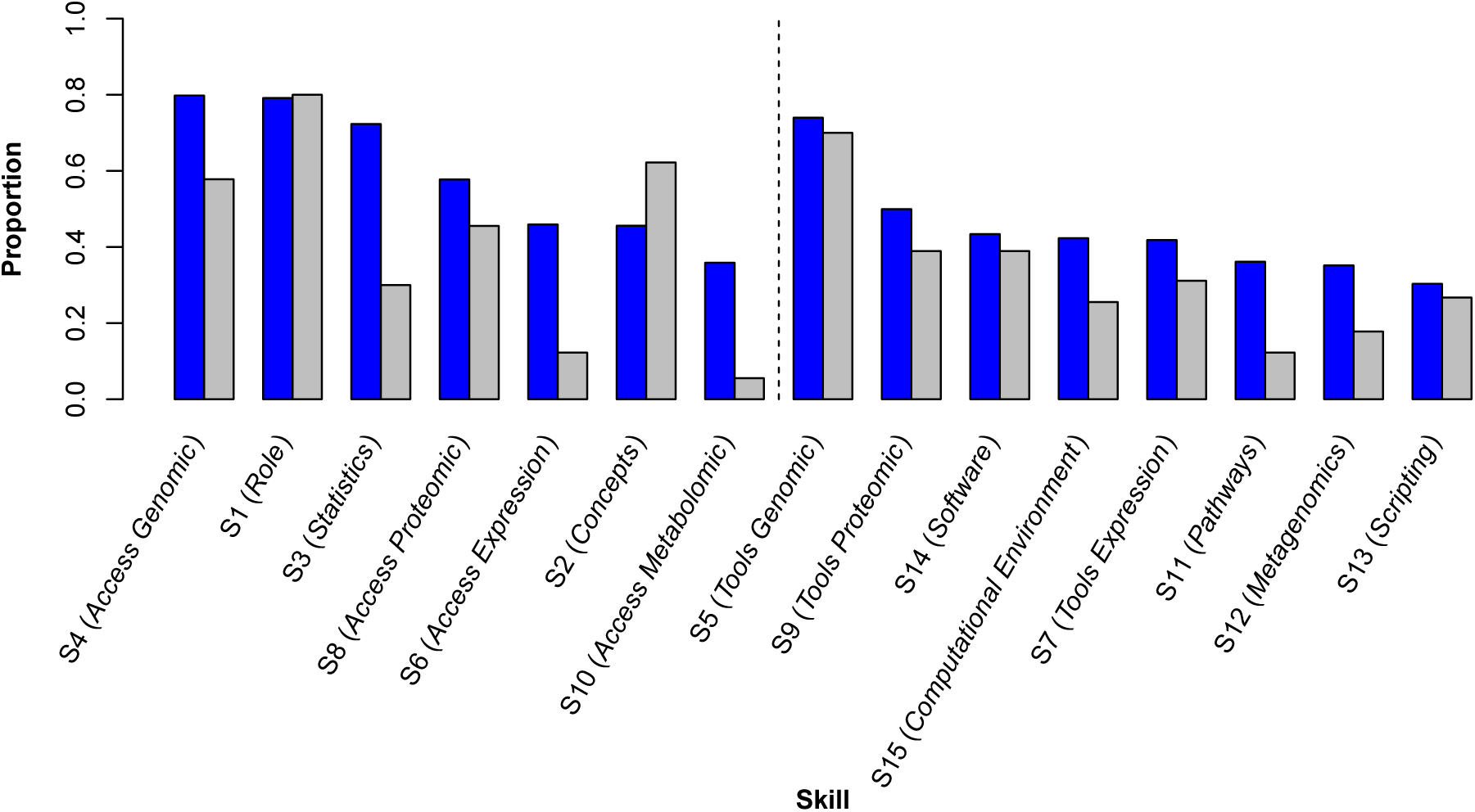
Comparison of the fifteen bioinformatics skills as rated by survey respondents versus coverage in the syllabi. Skills are shown with the proportion of survey responses rating the skill as either “Very Important” or “Extremely Important” (blue bars) and the proportion of submitted syllabi that exhibited evidence of the skill (grey bars). Skills requiring familiarity with a concept (“knowing” skills) are to the left of the vertical dashed line; skills requiring direct engagement (“practice” skills) are to the right. In their respective categories, skills are presented in order of decreasing proportion of survey responses rating the skill as Very or Extremely Important.

Syllabi also varied substantially in the number of survey skills covered. On average, a syllabus covered 5.5 skills (with a median of six skills addressed). Although all skills were covered in aggregate across the submitted syllabi, no syllabi covered more than thirteen out of the fifteen skills, suggesting the difficulty of covering all the skills in a single class (Fig. S2).

#### Core Competencies

The nine NIBLSE Core Competencies, developed using the iterative process described in Materials and Methods, are presented in Table 2. During this process, several skills that received lower ratings on the survey (S10, S11, S12) were dropped or combined into a more general competency such as C4 (“Use bioinformatics tools to examine complex biological problems”) and C6 (“Explore and/or model biological interactions, networks, and data integration using bioinformatics”). While scripting and use of the command line (S13) received relatively lower scores from survey respondents, the strong support for these emerging skills from the most recently trained respondents resulted in their inclusion in the final list. As mentioned above, the NIBLSE Core Competencies are intended to serve as a guide for institutions and departments as they integrate bioinformatics into their own life sciences curricula.

**Table 2.** The NIBLSE Bioinformatics Core Competencies for Undergraduate Life Scientists. The bioinformatics competencies that NIBLSE recommends undergraduate life sciences students have by the time they graduate. As discussed in the narrative, they are informed by the results of the national NIBLSE survey, analysis of ninety syllabi with bioinformatics content, and the cumulative expertise and experience of the authors. Following each competency is a list of three representative examples illustrating the competency.

|  |
| --- |
| **C1. Explain the role of computation and data mining in addressing hypothesis-driven and hypothesis-generating questions within the life sciences.** Life sciences students should have a clear understanding of the role computing and data mining play in modern biology. Given a traditional hypothesis-driven research question, students should have ideas about what types of data and software exist that could help them answer the question quickly and efficiently. They should also appreciate that mining large datasets can generate novel hypotheses to be tested in the lab or field. |
| • Compare and contrast computer-based research with wet-lab research.<br>• Explain the role of computation in finding genes, detecting the function of protein domains, and inferring protein function.<br>• Describe the role of various databases in identifying potential gene targets for drug development. |
| **C2. Summarize key computational concepts, such as algorithms and relational databases, and their applications in the life sciences.** To make use of sophisticated software and database tools, students should have a basic understanding of the principles upon which these tools are based and should be exposed to how these tools work. |
| • Explain the underlying algorithm(s) employed in sequence alignment (e.g., BLAST).<br>• Modify software parameters to achieve biologically meaningful results.<br>• Explain how data are organized in relational databases (e.g., NCBI and model organisms databases). |
| **C3. Apply statistical concepts used in bioinformatics.** In addition to the basic statistics found in many biology curricula, modern life scientists should have an understanding of the statistics of large datasets and multiple comparisons. |
| • Understand that there is a probability of finding a given sequence similarity score by chance (the $P$ value) and that the size of the target database affects the probability that you will see a particular score in a particular search (the E-value).<br>• Explain the statistical modeling used to identify differentially expressed genes.<br>• Interpret data from a well-designed drug trial. |
| **C4. Use bioinformatics tools to examine complex biological problems in evolution, information flow, and other important areas of biology.** This competency is written broadly so as to encompass a variety of problems that can be addressed using bioinformatics tools, such as understanding the evolutionary underpinnings of sequence comparison and homology detection; distinguishing between genomic sequences, RNA sequences, and protein sequences; and |
| interpreting phylogenetic trees. “Complex” biological problems require that students should be able to work through a problem with multiple steps, not just perform isolated tasks. |
| • Using multiple lines of evidence, annotate a gene.<br>• Develop and interpret a “tree of life” based on a BLAST search, multiple alignment, and phylogenetic-tree building.<br>• Explain how a mutation in a gene causes cancer, using a genome browser to identify the gene, transcript, and affected protein, and tools such as OMIM, GO, and KEGG to place it in the context of a function and pathway important to the disease. |
| **C5. Find, retrieve, and organize various types of biological data.** Given the numerous and varied datasets currently being generated from all of the ‘omics fields, students should develop the facility to identify appropriate data repositories, navigate and retrieve data from these databases, and organize data relevant to their area of study in flat files or small local stand-alone databases. |
| • Navigate and retrieve data from a genome browser.<br>• Retrieve data from protein and genome databases (e.g., PDB, UniProt, NCBI).<br>• Store and interrogate small datasets using spreadsheets or delimited text files. |
| **C6. Explore and/or model biological interactions, networks, and data integration using bioinformatics.** Modeling of biological systems at all levels, from cellular to ecological, is being facilitated by technological and algorithmic advances. These models provide novel insights into the perturbations in systems that can cause disease, interactions of microbes with various eukaryotic systems, how metabolic networks respond to environmental stresses, etc. Students should be familiar with the techniques used to generate these analyses and should be able to interpret the outputs and use the data to generate novel hypotheses. |
| • Predict impact of a gene knockout on a cell-signaling pathway.<br>• Analyze gene expression data to build an expression network.<br>• Analyze metagenomics data from microbial samples obtained from environmental sources. |
| **C7. Use command-line bioinformatics tools and write simple computer scripts.** Most biological datasets (e.g., genomic and proteomic sequences, BLAST results, RNASeq and resulting differential expression data) are available as text files; the most powerful and dynamic way to interact with these datasets is through the command line or shell scripting. Students should be able to manipulate their own data and to create and modify complex data processing and analysis workflows. |
| • Run BLAST using command line options.<br>• Build and run statistical analyses using environments such as R or Python scripts.<br>• Write simple shell scripts to manipulate files. |
| **C8. Describe and manage biological data types, structure, and reproducibility.** This competency addresses two distinct concerns: 1) each of the varied ‘omics fields produces data in formats particular to its needs, and these formats evolve with changes in technologies and refinements in |
| downstream software; and 2) all experimental data is subject to error and the user must be cognizant of the need to verify the reproducibility of their data. Students need to develop an awareness of, and ability to, manipulate different data types given the versioning of formats. They also need to exercise caution, to carry out appropriate statistical analyses on their data as part of normal operating procedures and report the uncertainty of their results, and to provide the relevant information to enable reproduction of their results. |
| ● Describe the various sequence formats used to store DNA and protein sequences (e.g., FASTA, FASTQ).<br>● Understand the representation of gene features using Gene Feature Format (GFF) files.<br>● Compare reproducibility of biological and technical replicate data (e.g., transcriptomic data) using statistical tests (Spearman rank test and false discovery calculations). |
| **C9. Interpret the ethical, legal, medical, and social implications of biological data.** The increasing scale and penetration of human genetic and genomic data has greatly enhanced our ability to identify disease-related loci, druggable targets, etc. and to identify potential genes for replacement with developing techniques. However, with this information also comes many ethical, legal, and social questions; suggested resolutions are often outpaced by the technological advances. As part of their scientific training, students should debate the medicinal, societal, and ethical implications of these information sets and techniques. |
| ● Explain the implications, good and bad, of being able to walk into a doctor's office and have your genome sequenced and analyzed or of being able to obtain information from direct-to-consumer testing services.<br>● Be able to discuss different perspectives about who should have access to this data and how it should be protected.<br>● Describe how the scientific community protects against the falsification or manipulation of large datasets. |

## Discussion

Faculty from a wide range of institutions showed strong support for greater integration of bioinformatics into undergraduate life sciences curricula: 95% of the 1,260 respondents agreed with the statement “I think bioinformatics should be integrated into undergraduate life sciences education.” However, there are some differences among faculty perspectives at different types of institutions. While most respondents value the ability to retrieve information from public databases and use existing software tools to analyze data, respondents at Doctoral institutions place a higher priority on computational skills, such as being able to operate in multiple computational environments and being able to write short programs. A possible explanation for this difference is that respondents at research-based institutions are more directly exposed to the necessity of using these computational skills on a day-to-day basis, whether by themselves, their students, or their colleagues. These findings provide insight into different educational perspectives and help us better understand what barriers institutions and departments may face as they integrate bioinformatics into their own life sciences curricula.

It is important to keep in mind the distinctions between the educational needs of bioinformaticians and life scientists, as well as the difference between the goals of undergraduate education and graduate or professional education. Up to this point, much of the discussion of bioinformatics education in the literature has focused on the education of bioinformaticians or on graduate or professional development. However, some authors have addressed the question of bioinformatics education for undergraduate life sciences students. In particular, attendees of the first and second Workshop on Education in Bioinformatics and Computational Biology in 2008 (Taipei, Taiwan) and 2009 (Singapore), held as part of the International Conference in Bioinformatics, attempted to identify a minimum skill set for the training of bioinformaticians and life scientists with informatics capabilities (8). A consensus list of five essential bioinformatics skills was reported. These skill sets overlap considerably with the NIBLSE Core Competencies, indicating that there is agreement about the skills that are necessary for modern life sciences students. However, the sample size of this effort was small (*n* = 56, including students) and the authors were attempting to find a consensus that could be applied internationally in widely differing contexts.

The NIBLSE Core Competencies include a computational competency, consistent with other reports (7,15,18,19). Although many life sciences programs are not currently equipped to provide training in basic programming and operating in a command-line environment, these are important skills that enable students to manipulate and analyze modern biological data. (As an added benefit, creating classes to teach these skills provides an opportunity for interdepartmental course development.) This does not mean that we recommend that life sciences students be able to write complex software applications or be trained to develop graphical user interfaces, but being able to write short programs and run command-line programs gives them flexibility in analyzing data and, perhaps more importantly, provides them with a better understanding of the data itself. The pedagogical literature from a variety of fields is clear that students learn more when they engage with data more deeply, as opposed to entering data into a “black box” and reporting the results (16,38–42). Thus, as with any laboratory technique—e.g., PCR, dissection, or microscopy—bioinformatically-literate undergraduates don’t need to be experts but should be expected to have basic skills in these areas when they graduate.

In addition to the survey data, the analysis of syllabi submitted by survey respondents shows that a variety of bioinformatics topics are already covered nationwide. The fact that bioinformatics can be performed relatively inexpensively with freely available data and software makes it an attractive way for students to engage in research experiences and inquiry-based learning (3,38,39). We would argue that this training would ideally occur in an integrated manner throughout a life sciences curriculum, as opposed to being isolated in a single course. Thus, there is no need to remove particular topics and replace them with “bioinformatics” units. Instead, we encourage faculty to find ways to incorporate bioinformatics techniques and applications as a way of exploring existing concepts in their curricula.

Among the syllabi analyzed, there appears to be a greater emphasis on “knowing” rather than “doing.” This gap may reflect the degree of instructor training: if an instructor has little training or experience with bioinformatics, it is understandably easier to introduce a technique or concept in a lecture than it is to develop and implement an in-depth exercise. This idea is supported by the fact that survey respondents frequently referred to the lack of available teaching resources in this area. One effort to address this problem was the creation of the Bioinformatics Learning Framework for *CourseSource*, which publishes evidence-based learning resources (31). The issue of training was also raised repeatedly in the open-ended comments on the survey, with many respondents indicating that they wanted to integrate more bioinformatics in their courses but felt they lacked the necessary training to do so. This training deficit is a long-term problem that will be difficult to address. On the one hand, a variety of resources are currently available for faculty to receive training in bioinformatics or to educate themselves to address the training need. On the other, many faculty in the survey indicated they needed more training. Thus, it’s not clear if the lack of training is due to faculty not being aware of the training resources available or if the existing resources are not useful to them because they cost too much, require too much time, are too advanced, or require too much work to adapt to their specific courses. Barriers to integrating bioinformatics into life sciences education, and our suggestions for overcoming them, are the topics of a separate paper, now in preparation. Note that NIBLSE was formed in large part to provide resources to faculty to help overcome these kinds of barriers.

To conclude, the analysis presented here provides evidence of strong, widespread agreement that undergraduate life sciences students need to be trained in bioinformatics, and it led to the development of a framework of topics for this training. Although the results presented here could potentially be skewed— those interested in this topic could have been more motivated to respond to the survey—given the large number of responses and the small margin of error in them (see Materials and Methods), we feel that we can take the results of the survey as accurately representing the opinions of the US life sciences education community as a whole. The goal of NIBLSE is to integrate existing efforts in bioinformatics education and to provide new resources to enable more faculty to infuse bioinformatics into their curricula. The NIBLSE Core Competencies presented here are expected to be a valuable guide to these efforts.

## Acknowledgements

This material is based upon work supported by the National Science Foundation under Grant Number 1539900 to A.G.R., E.W.T., E.D., W.M., and M.A.P. Any opinions, findings, and conclusions or recommendations expressed in this material are those of the authors and do not necessarily reflect the views of the National Science Foundation. The authors thank the members of the Genomics Education Partnership, Genome Solver, GCAT-SEEK, and NIBLSE networks for the feedback they provided during the development of the NIBLSE Core Competencies. The authors also thank Sarah Moulton of Moulton Editorial Services (Omaha, NE) for editing and formatting the final version of the manuscript.

## Author Contributions

The Core Competencies Team (M.A.W.S., C.H., M.S., and M.A.P.), analyzed the survey results and syllabi, came up with recommendations for the proposed list of final core competencies, and wrote the first draft of the manuscript.

The NIBLSE Core Competencies Working Group (M.A.W.S., C.H., M.S., S.R., A.G.R., T.M.S., E.W.T.,

L.R.W., and M.A.P) developed the survey and helped to distribute it.

The NIBLSE Leadership Team (A.G.R., E.W.T., E.D., W.M., and M.A.P.) organized the NIBLSE conference and decided on the final wording of the competencies.

All authors contributed to the development of the NIBLSE Core Competencies and to the manuscript.

